# Magnitude processing and integration entail perceptual processes independent from the task

**DOI:** 10.1101/2024.04.29.591641

**Authors:** Irene Togoli, Olivier Collignon, Domenica Bueti, Michele Fornaciai

## Abstract

The magnitude dimensions of visual stimuli, such as their numerosity, duration, and size, are intrinsically linked, leading to mutual interactions across them. However, it remains debated whether such interaction across dimensions, or “magnitude integration” effects, arise from low-level perceptual processes that are independent from the task performed, or whether they instead arise from high-level decision-making processes. We address this question with two experiments in which participants watched a series of dot-array stimuli modulated in numerosity, duration, and item size. In experiment 1 (task condition), the task required participants to either judge the numerosity, duration, or size of each stimulus. In experiment 2 (passive condition), instead, a separate group of participants passively watched the stimuli. The behavioral results obtained in the task show robust magnitude integration effects across all three dimensions. Then, we identify a neural signature of magnitude integration by showing that event-related potentials at several latency windows (starting at ∼100-200 ms after stimulus onset) can predict the effect measured behaviorally. In the passive condition, we demonstrate an almost identical modulation of brain responses, occurring at the same processing stages as during the task. Importantly, using a cross-condition multivariate decoding analysis, we demonstrate that brain responses to magnitude in the task condition can predict the response in the passive condition at specific latency windows. These results thus suggest that magnitude processing and integration likely occurs via automatic perceptual processes that are engaged irrespective of the task-relevance of the stimuli, and independently from decision making.

## INTRODUCTION

Magnitude dimensions such as numerosity, time, and space represent fundamental properties of the external world, as each of these dimensions provides essential information to understand and navigate the environment. Indeed, the perception of magnitudes organize our thoughts and experience by allowing us to appreciate how many objects are around us, their size and their spatial relations, and the duration and timing of the external events. While these dimensions are important in their own rights and studied in separate lines of research, a particularly interesting phenomenon is their integration and interaction, likely grounded on their shared computational structure (e.g., Walsh, 2003). Different magnitude dimensions seem indeed linked in a way that the perception of one dimension depends on the others, usually leading to mutual biases. For instance, a large object or a numerous set of items is perceived as lasting longer in time compared to a smaller object or fewer items (e.g., Xuan et al., 2007). Vice versa, a longer stimulus may appear bigger or more numerous than a shorter one (e.g., Javadi & Aichelburg, 2012; Lambrechts et al., 2013; Togoli et al., 2021).

These mutual influences across magnitude dimensions – or “magnitude integration” effects – represent one of the core phenomena characterizing magnitude perception, and have inspired important theories like the “a theory of magnitude” (ATOM) framework (Walsh, 2003). According to ATOM, the processing of different magnitudes culminates in a generalized magnitude system encoding different dimensions with the same neural code. This in turn would allow the interaction of magnitude information in the service of perception and behavior. This view has been however challenged by the idea that biases across magnitudes, and especially space and time, may stem from the linguistic labels assigned to them, and how we conceptualize these dimensions at a linguistic rather than at a perceptual level (“metaphoric theory;” e.g., Casasanto & Boroditsky, 2008). While evidence has now been accumulated against a purely linguistic/conceptual view of magnitude integration (Cai & Connell, 2015; Togoli, Bueti, et al., 2022; Whitaker et al., 2022), other theories have proposed that magnitudes interact at a more cognitive rather than perceptual level, as a working memory interference (Cai et al., 2018), or as a response bias (Yates et al., 2012). Namely, according to these ideas, the interference across magnitudes would occur either because of different memory traces nudging each other while stored in working memory, or because of a bias in the response selection due to the similar response codes of different magnitudes (i.e., “more” vs. “less”). In both these cases, the interference would not affect how magnitudes are perceived, but only their memory traces or the way they are judged. Finally, based on neuroimaging data, it has been recently proposed (Hendrikx et al., 2024; Tsouli et al., 2022) that the interaction could arise from the processing of different dimensions in partially overlapping cortical maps, but without involving a common neural code (Fortunato et al., 2023; Harvey et al., 2013, 2015; Hendrikx et al., 2022, 2024; Protopapa et al., 2019).

At which processing stage magnitude integration arises thus remains debated. Mixed evidence indeed seems to support both the “low-level,” perceptual interpretation, and the “high-level” interpretation based on memory and/or decision making. For example, results from Cai et al. (2018) show that the duration judgements can be biased by the length of a stimulus only when the length information is provided before the duration judgment has started, suggesting that the bias occurs as an interference between memory traces. Furthermore, electroencephalographic (EEG) evidence from Cui et al. (2022) shows that the interference of length on duration is reflected by event-related potentials (ERPs) usually associated with working memory (i.e., the P2 and P3b component). Conversely, other results show that integration effects do not occur every time two magnitudes are presented, as one would expect for instance from a response bias, but only when the two dimensions are conveyed by the same stimulus (e.g., a dot array with a given numerosity flashed to mark the onset and offset of a duration; Togoli, Bueti, et al., 2022). Instead, when two dimensions like duration and numerosity are conveyed by different stimuli (i.e., a texture marking the onset and offset of a duration, flashed on top of a dot array), the effect reverses becoming repulsive (i.e., the more numerous the stimulus is, the shorter it is perceived to last). This suggests that magnitude integration effects are not the result of a simple interference between different types of information, but involve perceptual binding processes.

To further assess the nature of the magnitude integration phenomenon, here we compare the neural (EEG) signature of magnitude integration when magnitudes are actively judged in a task, versus when they are passively watched. In the first experiment, the participants judged either the numerosity, the duration, or the item size of dot-array stimuli against a reference presented before the start of each block (task condition). In a second experiment, a separate group of participants passively watched a similar stream of dot-array stimuli modulated in numerosity, duration, and item size (passive condition). Our hypothesis is that if magnitude processing and integration entail perceptual processes, then similar magnitude-sensitive neural signatures should be observed irrespective of whether the participants are actively judging magnitudes or not. Conversely, if the integration effect hinges upon memory or decision-making, then brain activity during the task should show a unique neural signature not generalizing to the passive condition, where the magnitudes are neither memorized nor judged. To test this hypothesis, we first identify a neural signature of magnitude integration in the task condition, by assessing the extent to which the brain responses could predict the integration effect measured behaviorally. We then compare such a neural signature with brain activity evoked by the different magnitudes in the passive condition. Finally, to achieve a quantitative measure of how similar the brain responses in the two conditions are, we use a multivariate cross-condition “decoding” analysis. With this analysis, we thus assess the extent to which magnitude-sensitive brain responses during passive viewing can be predicted from the data of the task condition. If magnitude processing and integration entail similar mechanisms engaged irrespective of the task, then the brain responses to the magnitudes in the passive condition should be decodable based on the task data. Otherwise, if the task engages specific mechanisms resulting in different patterns of brain activity, no cross-condition decoding should be observed.

## MATERIALS AND METHODS

### Participants

A total of 51 participants were tested in the study, with 20 participants tested in the task condition (13 females; age ± SD = 24.95 ± 4.21) and 31 separate participants tested in the passive condition (19 females; age ± SD = 23.96 ± 3.73). Two participants were excluded from data analysis in the passive condition due to corrupted EEG data files, leaving 29 participants included in the final analysis. Subjects were compensated for their participation in the study with 20 Euros. All participants read and signed a written informed consent form before the start of the session. All participants had normal or corrected-to-normal vision, and reported no history of neurological, psychiatric, or developmental disorder. The study was approved by the ethics committee of the International School for Advanced Studies (Protocol 10035-III/13), and was designed to be in line with the Declaration of Helsinki. The sample size tested in the task condition was determined a priori based on the expected magnitude integration effect as observed in previous studies. Specifically, we took the average of the lowest effect sizes of the magnitude integration effects observed in Togoli et al. (2021) (i.e., average of effects at the smallest magnitude levels in the time and numerosity task of Exp. 1a; d = 0.87), the effect size observed in Togoli, Bueti, et al., (2022) (average of effects in Exp. 2; d = 0.74), and in Togoli, Fornaciai, et al. (2022) (i.e., average of effects at the intermediate magnitude levels in the time and numerosity task; d = 1.21). The resulting average effect size was d = 0.94. Considering a power of 95% and a two-tailed distribution, a power analysis indicated a total estimated sample size of 17 participants, which we rounded up to 20.

Since a measure of the behavioral effect cannot be obtained in the passive viewing paradigm, the sample size in the passive condition was set to be similar to a series of previous EEG studies in numerosity perception, which included on average 25-30 participants (Fornaciai et al., 2017; Fornaciai & Park, 2017, 2018, 2020, 2021).

### Apparatus and stimuli

The stimuli used in both conditions were arrays of black and white dots (50%/50% proportion), presented on a grey background at maximum contrast. The stimuli were generated using the routines of the Psychophysics Toolbox (v.3; Kleiner M et al., 2007; Pelli, 1997) in Matlab (r2021b, The Mathworks, Inc.), and presented on a 1920×1080 LCD monitor running at 120 Hz, which encompassed a visual angle of 48×30 deg from a viewing distance of 57 cm. The dot-array stimuli were generated online in each trial, with the dots scattered pseudo-randomly within a circular aperture with a variable radius spanning from 200 to 400 pixels (pseudo-randomly determined in each trial). In the task condition, the dot-array stimuli could have a numerosity of 8, 12, 16, 24, or 32 dots, a duration of 100, 140, 200, 280, or 400 ms, and item size (i.e., the radius of each item in the array) of 3, 4, 6, 8, or 10 pixels, for a total of 125 unique combinations of the three magnitudes. The different ranges were designed to be approximately spaced in a Log2 scale. The reference stimulus that the participants used as a comparison (presented at the beginning of the session and before each block) had the middle value of the three ranges (16 dots, 200 ms, 6 pixels). The stimuli in the passive condition had smaller magnitude ranges to make the modulation of magnitudes subtler and less obvious, in order better mask the true aim of the experiment. Namely, the stimuli could have a numerosity of 12, 16, or 24 dots, a duration of 140, 200, or 280 ms, and item size of 4, 6, or 8 pixels. Again, these ranges were designed to be approximately spaced in a Log2 scale. In both conditions, the stimuli were presented at the center of the screen.

### Procedure

*Task condition*. In the task condition, participants performed a magnitude classification task, with the dimension judged in each trial determined by a retrospective cue (i.e., presented after the offset of the stimulus). First, participants watched a reference stimulus representing the middle of the magnitude ranges used in the experiment and were instructed to remember it and judge the stimuli in main sequence based on it. The reference was presented 10 times (randomizing the positions of the dots) before the start of the session and repeated 5 more time before the start of each block of trials. In the session, participants were instructed to keep their gaze on a central fixation point. Each trial started with the presentation of the fixation cross (a “X” at the center of the screen). After 750 ms, the dot-array stimulus was presented replacing the fixation cross, and was displayed for 100-400 ms according to the duration selected in the trial. After an interval of 600 ms from the offset of the stimulus, the retrospective cue was presented at center of the screen. The cue could be either “N,” “T,” or “S,” respectively instructing the participant to judge the numerosity, duration (i.e., time), or item size of the stimulus. The cue remained on the screen for 600 ms. After that, the cue was replaced by a red X (fixation cross), instructing the participant to provide a response. According to the cue, the participant was asked to indicate whether the stimulus had a higher or lower numerosity, a longer or shorter duration, or a bigger or smaller item size compared to the reference. The response was provided by pressing either the down arrow (lower/shorter/smaller) or the up arrow (higher/longer/bigger) on a standard computer keyboard. After providing a response, the next trial started after a variable inter-trial interval of 500 ± 50 ms. Each participant completed a total of 1,250 trials (10 blocks of 125 trials), corresponding to a total of 10 repetition of each unique combination of numerosity, duration, and dot size. The three tasks were randomly intermixed within the same blocks. No feedback was provided to participants about their response.

*Passive condition*. In the passive condition, the participants watched a series of stimuli modulated in numerosity, duration, and size. Each stimulus was presented centrally on the screen, and successive stimuli were separated by an inter-stimulus interval of 980 ± 50 ms. In order to make participants attend the stream of stimuli, they were asked to detect occasional oddball stimuli defined by a reduced contrast compared to the rest of the stimuli (oddball detection task). The oddball stimuli represented 3.7% of the total stimuli presented. Participants were thus instructed to press the space bar on the keyboard as fast as they could once they detected an oddball stimulus. This occasional simple detection task was designed to avoid drawing attention to any of the magnitude dimensions of the stimuli, while encouraging the participants to watch the stimuli. On average, the detection rates (± SD) of the oddball were 93% ± 1.3%, and the average reaction time was 313 ± 11 ms. In the passive condition, participants completed a total of 2,160 trials (8 blocks of 270 trials), for a total of 80 repetitions of each combination of stimulus magnitudes. The higher number of trials tested in the passive condition compared to the task condition was chosen to compensate for the smaller magnitude ranges used, i.e., in order to ensure that we could measure robust brain responses to the magnitudes also in this condition. Overall, participants were only instructed to watch the stream of stimuli and respond to the oddball, and the magnitudes were never mentioned in the instructions and recruiting materials.

### Behavioral data analysis

In the task condition, the magnitude judgement performance and the integration effect were assessed by first computing the point of subjective equality (PSE). The PSE reflects the accuracy in the task, and the perceived magnitude of the stimuli compared to the reference magnitude. Data reflecting each specific task (i.e., all the trials in which a specific cue was presented) were used to compute the proportion of “more” (numerosity task), “longer” (duration task), or “bigger” (size task) responses as a function of both the task-relevant magnitude and the other (interfering) magnitudes. A psychometric (cumulative Gaussian) function was then fitted to the distribution of proportion of response, according to the maximum likelihood method (Watson, 1979). Specifically, within each task, the psychometric function was fitted separately for each level of each of the other “interfering” magnitudes, in order to assess the difference in PSE due to the task-irrelevant magnitudes. For example, when analyzing the performance in the numerosity task, the fit was performed separately for each level of duration and each level of item size. To account for errors unrelated to the magnitude of the stimuli and lapses of attention, a finger-error rate correction of 2.5% (Wichmann & Hill, 2001) was applied. This correction reduces the asymptotic levels of the fit by a proportion corresponding to the rate, in order to account for the random errors preventing the proportion of responses to converge to 0% and 100% at the lower and higher end of the range, respectively. The PSE was computed as the numerosity/duration/size level corresponding to chance level (50%) responses (i.e., the median of the psychometric curve). This procedure allowed us to compute individual measures of PSE for the different combinations of task relevant and interfering dimensions (e.g., numerosity PSE when duration was 100 ms, 140 ms, 200 ms, 280 ms, and 400 ms, and the same for size, and similarly for the other tasks). When performing the fit according to a given task-relevant and interfering magnitude, the other dimension was collapsed. Doing so, each data point in the fitting procedure represented the average of 25 repetitions of the specific combination of task-relevant and interfering dimension. Additionally, we assessed the precision in the task in terms of the Weber fraction (WF). The WF was computed as the ratio between the just noticeable difference (JND; the slope of the psychometric curve) and the PSE. To assess the difference in the average WF across the three types of tasks, we used a one-way repeated measures ANOVA.

In order to better define the effect of the interfering magnitudes in each task, we then used the PSE to compute a “magnitude integration effect” index according to the following formula:

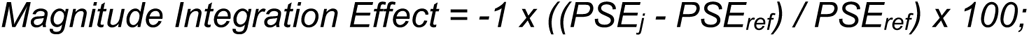

Where *PSEref* correspond to the PSE obtained when the interfering magnitude considered was the same as the reference, and *PSEj* to the PSE corresponding to each other level of the interfering magnitude (either lower or higher than the reference). The change in sign (-1) was added in order to make the interpretation of the index easier. Namely, doing so a positive index means that the task-relevant magnitude is overestimated, while a negative index means that the magnitude is underestimated. To assess the significance of magnitude integration effects, we performed a series of linear mixed-effect model tests, assessing the integration biases on each type of judgement. Namely, we entered the magnitude integration effect as dependent variable, the ratio of each level of the interfering magnitude with the reference value and the magnitude itself (i.e., “numerosity,” “duration,” or “size”) as predictors, and the subject as the random effect (Magnitude integration effect ∼ Ratio x Magnitude + (1|subj.)). Interactions found between ratio and magnitude were followed up with additional LME tests within each interfering magnitude dimension. The LME models were chosen in this case (i.e., instead of ANOVAs) as the ratio is a continuous variable.

Finally, in order to assess the relationship between ERPs and the behavioral effect, we computed a measure of ΔPSE. This measure reflected the difference in PSE between each level of the interfering dimension and the middle magnitude level corresponding to the reference. This was done to have a similar measure that can be related to the neural effect of different magnitudes (see below for more information about the ERP analysis). All the analyses and statistical tests on behavioral data were performed in Matlab (version r2021b).

### Electrophysiological recording and pre-processing

In both the task and passive condition, the EEG was recorded throughout the experimental session. EEG recording was performed by using the Biosemi ActiveTwo system (at 2048 Hz sampling rate), and a 32-channel cap based on the 10-20 system layout. To better monitor artifacts due to eye blinks and movements, we recorded the electro-oculogram (EOG) by means of a channel attached below the left eye of the subject. During the recording, we made sure to keep the electrode offsets as low as possible. Usually, electrode offset values were kept below 20 µV, but occasionally values up to 30 µV were tolerated.

The pre-processing of EEG data was performed offline in Matlab (version R2021b), using the functions of the EEGLAB (Delorme & Makeig, 2004) and ERPlab (Lopez-Calderon & Luck, 2014) toolboxes. In both conditions, the pre-processing involved the binning and epoching of data according to each unique combination of the different magnitudes. In the task condition, the binning was also performed separately for the three cues determining the specific task in each trial. In both conditions, epochs spanned from -200 to 700 ms, time-locked to the onset of each stimulus. In the case of duration, ERPs were later re-aligned to the offset of the stimuli. This was done as we expected an effect of duration only after the presentation of the stimuli had fully unfolded.

In both conditions, after the epoching, the EEG data was band-pass filtered with cut-offs at 0.1 and 40 Hz. Moreover, to clean up the data as much as possible from artifacts such as eye movements and blinks, we performed an independent component analysis (ICA) aimed at removing identifiable artifacts and other potential sources of systematic noise. After the ICA, we additionally applied a step-like artifact rejection procedure, with an amplitude threshold of 40 μV, a window of 400 ms, and a step size of 20 ms. This was done in order to further remove any remaining large artifact from the EEG data. On average, this led to the exclusion of 2.26% ± 1.73% of the trials in the task condition, and 0.6% ± 1.2% in the passive condition. Finally, we computed the event-related potentials (ERPs) by averaging EEG epochs within each bin, and further low-pass filtered the signal with a 30-Hz cut-off.

### Event-related potentials analysis

In both conditions, the ERP analysis was performed by considering the average of the same set of three occipito-parietal target channels, selected a priori based on previous studies (Fornaciai et al., 2017; Fornaciai et al., 2023; Tonoyan et al., 2022). The chosen target channels were Oz, O1 and O2.

First, we plotted the ERPs corresponding to each magnitude and computed the linear contrast of ERPs. The weights of the linear contrast computation were [-2 -1 0 -1 -2] in the task condition, and [-1 0 1] in the passive condition, based on the number of levels of each magnitude tested in the two conditions. Moreover, we computed an additional measure of contrast by computing the difference in ERP amplitude for each level of each magnitude and the two extreme levels of the other two interfering dimensions. For example, for each level of numerosity we computed the difference in amplitude between the extreme levels of duration and size, and so on for the other dimensions. We then averaged this measure across the different levels of each magnitude. To assess the significance of the magnitude contrasts, we performed a series of one-sample t-tests against zero. To control for multiple comparisons, we applied a false discovery rate correction with q = 0.05. Finally, we computed a measure of ΔERP as the difference in amplitude between the ERPs corresponding to the middle level of the range (corresponding to the reference in the task condition) and each other level of the magnitude ranges. In the task condition, the ΔERP was computed separately according to the task. Doing so, we thus computed a measure of the effect of numerosity in the size and duration task, and so on for the other magnitudes. This measure was computed to have an index of the effect of magnitudes on ERPs similar to the ΔPSE computed from the behavioral results, in order to more easily relate the ERP and behavioral measures of the effect in the analysis of the task condition. To assess the significance of the ΔERP, in the task condition we performed a series of LME tests, entering ΔERP as the dependent variable, the ratio of each magnitude level and the middle level as predictor, and the subjects as the random effect (ΔERP ∼ Ratio + (1|subj)). In the passive condition, we instead performed a series of paired t-tests (i.e., due to the smaller number of levels of each magnitude range). In both cases, the tests were performed on small 10-ms (step = 5 ms) windows in a sliding-window fashion, and the significance was corrected with FDR (q = 0.05). Additionally, we considered significant only clusters of consecutive significant time-points larger than 10 ms (i.e., three consecutive significant tests or more).

Only in the task condition, we further assessed the relationship between ΔERP (i.e., neural effect of magnitude) and ΔPSE (i.e., behavioral effect of magnitude) via a series of LME tests. In this case, we entered the ΔPSE as the dependent variable, the ΔERP as the predictor, and the subjects as the random effect (ΔPSE ∼ ΔERP + (1|subj)). This analysis was performed again across a series of 10-ms windows with step = 5 ms. Finally, the analysis was restricted to the latency windows showing a significant modulation of ΔERP by the magnitudes (see above), and we considered statistically significant only clusters of at least three consecutive significant windows. This analysis was aimed at assessing the extent to which magnitude-sensitive brain responses can predict the effect measured behaviorally.

### Multivariate analysis

We relied on a multivariate approach in order to more directly compare the brain activity related to magnitude processing in the two conditions. Namely, we assessed the extent to which training a classifier on the data (ΔERP) from the task condition allows to decode magnitude-sensitive activity in the passive condition. First, to create the training dataset, we computed for each participant an average measure of ΔERP in the task condition, based on the difference between the amplitude relative to lower (8, 12 dots; 100, 140 ms; 3, 4 pixel) and higher (24, 32 dots; 280, 400 ms; 8, 10 pixel) magnitude levels and the amplitude of the middle magnitude level (16 dots, 200 ms, 6 pixel). Thus, the training dataset was composed by a number of data points corresponding to the number of participants in the task condition (N = 20), with each data point corresponding to the average data of a single participant (i.e., one data point for each specific class entered into the analysis). The same was done for each participant in the passive condition group to create the test dataset, based on the specific levels of the magnitudes used in this condition (12 and 24 dots, 140 and 280 ms, 4 and 8 pixels; respectively for the magnitudes lower and higher than the middle value). Again, this procedure resulted in a set of data points (N = 29) each corresponding to the average data of each participant, one for each of the two classes entered into the analysis (i.e., magnitudes “lower” or “higher” than the middle value, with different dimensions tested separately in different iterations of the procedure). In the decoding procedure, a linear classifier (support vector machine, C = 1) was trained on the task condition dataset, according to a leave-two-out cross-validation procedure. Namely, a datapoint for each of the two levels of magnitude (“lower” and “higher”) was left out from the training set. To test the classifier, we used the two data points from a single participant in the passive condition group. Thus, in each iteration of the analysis, the training was performed on 19 + 19 data points (corresponding to the two classes being decoded) based on the task condition data, and the testing on 1 + 1 data points (again corresponding to the two classes) from the passive condition dataset. This procedure was repeated to include all the combinations of training data, iteratively leaving out each data point from the training set and testing the classifier again with the same test data points, and averaging the classification accuracies obtained in the different cross-validation steps. The analysis was repeated considering each set of test data (i.e., the data from each participant in the passive condition) separately. This in turn resulted in a distribution of classification accuracy values corresponding to the number of participants in the passive condition group. Moreover, the analysis was performed independently across a series of small time windows (15 ms width, 5 ms step; i.e., to increase the signal-to-noise ratio), throughout the epoch. Finally, the training and testing procedure was performed by considering a set of five occipital and parieto-occipital channels (PO3, O1, Oz, O2, PO4). To test for the significance of the distribution of classification accuracies obtained at each time window, we used a permutation (sign flipping) procedure. Namely, at each time window, we subtracted 0.5 from the classification accuracies (i.e., the chance level), and swapped the sign of half the values. This procedure was repeated 10,000 times taking random splits of the data, and we assessed the number of times that the classification accuracy of the sign-swapped data was equal or higher than the actual average classification accuracy observed. The proportion of times in which the simulated accuracy equaled or exceeded the actual value was considered the p-value of the test. The alpha level applied to these tests was 0.05. After the tests, we also applied a threshold of at least three consecutive time windows (i.e., only clusters of at least three consecutive time windows were considered significant). Note that we chose to use the passive condition data as the testing dataset in order to have a larger distribution of classification accuracy values to test with permutations, with the rationale of achieving more robust and stable results. We did not perform the analysis in the opposite direction (training with the passive data and testing with the task data) since the two sets of results would be difficult to average. Namely, due to the nature of this procedure, analyses in different directions would results in different distributions of classification accuracy values (29 when testing with the passive data, 20 when testing with the task data), making it difficult to combine them. All the analyses and statistical tests involving EEG data were performed in Matlab (version r2021b).

## RESULTS

In this study we measured the neural (EEG) signature of magnitude processing and integration with subjects either engaged in actively judging the magnitude of the stimuli, or passively watching the stimuli. Doing so, we aimed at comparing such signatures to better understand the nature of the magnitude integration phenomenon. Specifically, if magnitude integration depends on post-perceptual cognitive processes, we predicted to observe a unique signature of magnitude processing only when performing a magnitude task. Conversely, if the integration effect arises from automatic perceptual processes, then similar signatures of magnitude processing should be observed in both the task and during passive viewing.

### Task condition

In the task condition, participants performed a magnitude classification task of the numerosity, duration, and item size of dot-array stimuli. Which dimension to judge was indicated to the participants via a retrospective cue (in a trial-by-trial fashion), thus forcing them to attend the stimulus as a whole rather than focusing on a single dimension. The procedure of the task condition is depicted in Fig. 1A.

**FIGURE 1.**
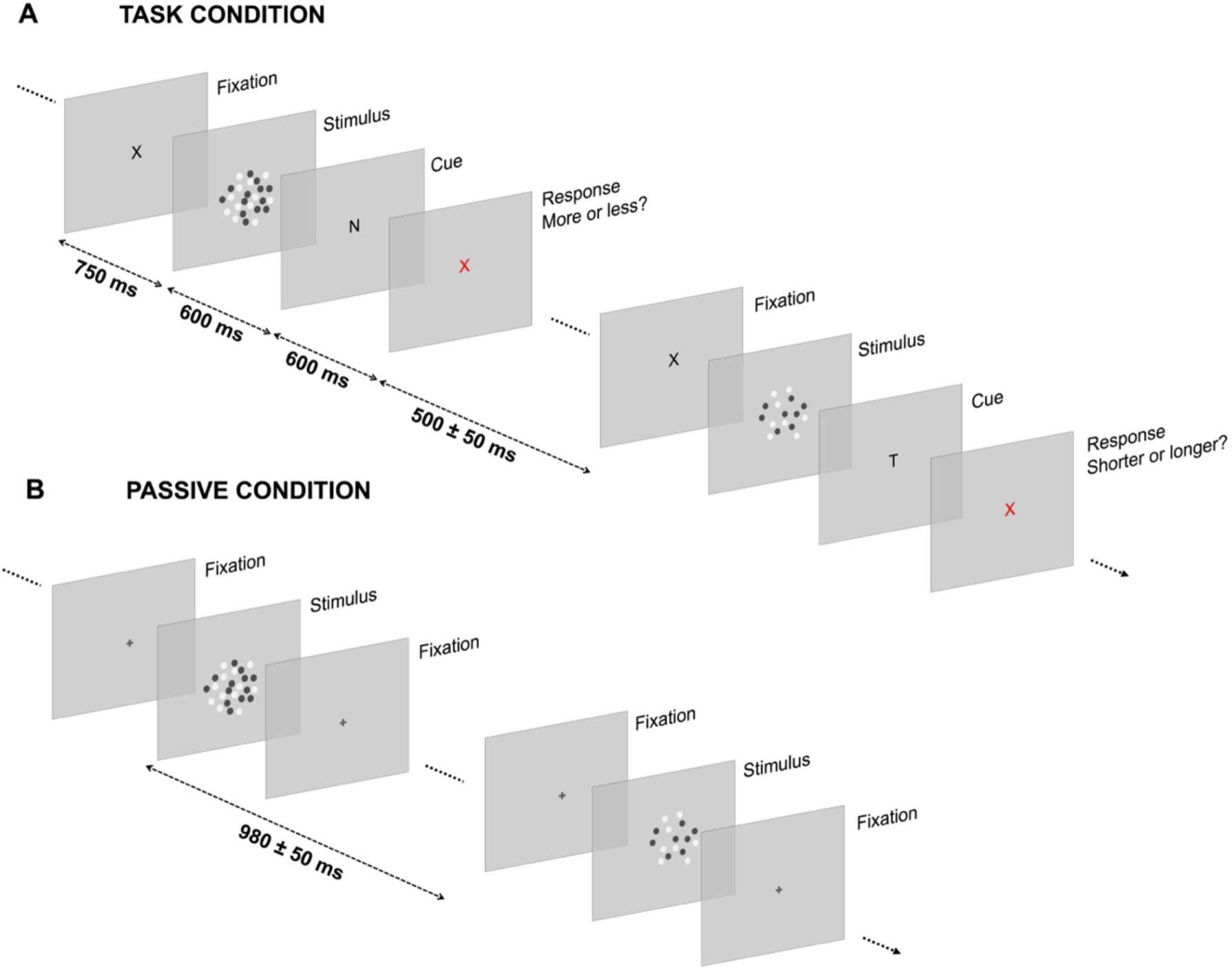
Experimental procedure. (A) Procedure of the task condition. While participants kept their gaze at the center of the screen (on a “X” that served as fixation cross), a stimulus was presented in each trial. The stimulus was modulated in numerosity (8-32 dots), duration (100-400 ms), and item size (i.e., the size of each item in the array; 3-10 pixels). After an interval of 600 ms from the offset of the stimulus, a cue appeared at the center of the screen indicating which stimulus dimension the participant had to judge. Namely, the cue could be a “N” (numerosity judgment), a “T” (i.e., duration judgement), or a “S” (size judgement). The cue remained on the screen for 600 ms, after which the fixation cross turned red indicating to provide a response. The participants were then asked to report whether the magnitude dimension indicated by the cue was bigger or smaller compared to a reference corresponding to the middle of the magnitude ranges (presented before the session and before each block of trials). After providing a response, the next trial started after a variable inter-trial interval (500 ± 50 ms). (B) Passive viewing condition. In the passive condition, participants watched a stream of dot-array stimuli modulated in numerosity (12-24 dots), duration (140-280 ms), and item size (4-8 pixels), while keeping their gaze on a central fixation cross. To ensure that participants watched the stimuli, they were asked to detect an occasional oddball stimulus (i.e., a dot array with reduced contrast) presented on 3.7% of the trials. Each stimulus was separated by a variable inter-stimulus interval of 980 ± 50 ms. Stimuli are not depicted in scale.

First, we assessed the behavioral effects of magnitude integration, which are shown in Fig. 2A-C. To assess the mutual biases across the different dimensions, we first computed the point of subjective equality (PSE; see *Behavioral data analysis*) as a measure of accuracy in the task. Then, we derived a magnitude integration effect index based on the difference in PSE caused by each level of the interfering magnitudes compared to the reference magnitudes (i.e., the middle levels of numerosity, duration, and size). To assess the significance of magnitude integration, we performed a series of linear mixed-effect (LME) regression models within each type of task. In the model, we entered the magnitude integration effects as the dependent variable, the ratio of each interfering magnitude level with the reference level, and the magnitude itself (e.g., “duration” and “size” in the numerosity task), as predictors, and the subjects as the random effect. The ratio (instead of the magnitude) was chosen as a predictor to test the effect of both interfering dimensions on each type of judgment within the same test.

**FIGURE 2.**
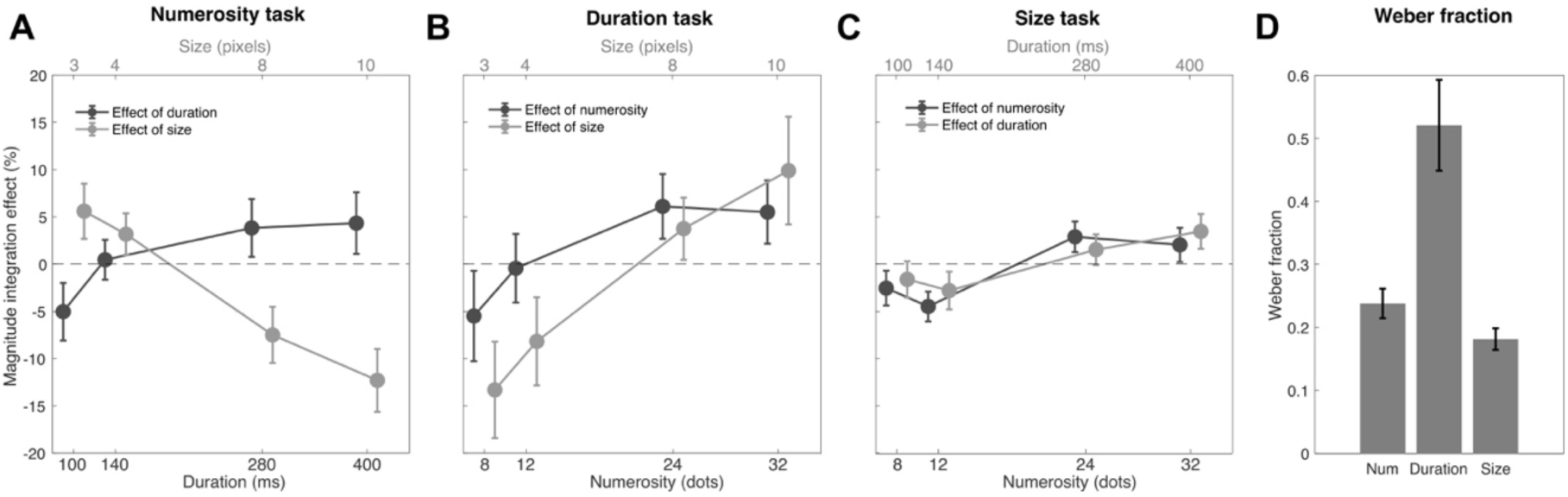
Behavioral effects of magnitude integration. (A) Magnitude integration effects of duration and size on numerosity judgements. The values of the two interfering dimensions are reported on the upper and lower x axes. (B) Magnitude integration effects on duration judgements. (C) Magnitude integration effect on size judgements. The data points are slightly jittered for the ease of visualization. (D) Average Weber fraction in the three tasks. Error bars are SEM.

In the numerosity task (Fig. 2A), we observed robust effects of both duration and size, although in opposite directions. While duration had a congruent effect (i.e., the longer the duration, the higher the perceived numerosity), size had an opposite, repulsive effect: the bigger the size of the dots, the lower the perceived numerosity. The LME test (adjusted-R^2^ = 0.22) indeed showed a significant interaction between ratio and magnitude (β = 20.96, t = 5.90, p < 0.001). This interaction was followed up with simpler LME tests considering each interfering dimension separately. The results of these additional tests showed that both duration (adj-R^2^ = 0.56, β = 5.47, t = 3.23, p = 0.002) and size (adj-R^2^ = 0.47, β = -15.50, t = -6.16, p < 0.001) induced significant biases on numerosity judgements. In the duration task (Fig. 2B), we observed again a significant interaction between ratio and magnitude (adj-R^2^ = 0.37, β = -12.02, t = - 2.48, p = 0.019), this time suggesting that size had a stronger influence on duration compared to the effect of numerosity on duration. Two follow-up LME tests however showed significant congruent biases induced by both numerosity (adj-R^2^ = 0.33, β = 7.47, t = 2.75, p = 0.007) and size (adj-R^2^ = 0.46, β = 19.30, t = 4.88, p < 0.001). Finally, looking at Fig. 2C it is clear that size was the magnitude most resistant to biases from other dimensions. Although weaker, the LME test (adj-R^2^ = 0.27) showed a significant main effect of ratio (β = 3.99, t = 3.02, p = 0.003), no main effect of magnitude (β = -1.31, t = -0.53, p = 0.59), and no interaction (β = 0.41, t = 0.22, p = 0.82). Overall, the behavioral results of the classification task showed systematic mutual biases across all the dimensions tested, albeit with some partial asymmetries.

To assess the participants’ precision in the task, we considered the Weber fraction (WF; computed as the ratio of the just noticeable difference and the PSE), which is shown in Fig. 2D. On average, size showed the lowest WF (0.18 ± 0.08), suggesting the highest precision in the task, followed by numerosity (0.24 ± 0.10), and finally duration (0.52 ± 0.32), which was the most difficult dimension to judge. A one-way repeated measure ANOVA (with factor “task”) confirmed that the WFs across the three types of task are significantly different (F(2,38) = 18.59, p < 0.001), in line with previous studies (Fornaciai et al., 2023).

After assessing the behavioral effects, we went on and addressed the neural signature of magnitude integration. First, we plotted the event-related potentials (ERPs) evoked by the different levels of each magnitude, irrespective of the task performed. The ERPs are shown in Fig. 3A-D. In the case of numerosity (Fig. 3A), we observed a first negative peak of numerosity-sensitive responses at 150-200 ms after stimulus onset, followed by weaker but more sustained responses throughout the epoch. In the case of duration, aligning the waves to the offset of the stimuli created a misalignment of the onset responses, which introduced a few spurious effects. In other words, a seemingly large deflection in the contrast amplitude is actually driven by a peak in amplitude of a single level of duration, rather than a consistent peak present at all levels of duration. However, we also observed a large modulation at around 300 ms after stimulus offset, that seems to genuinely reflect duration, as it involves a deflection evident at all three levels of duration. Finally, brain responses sensitive to item size showed the main peak at around 250 ms, with a large positive deflection.

**FIGURE 3.**
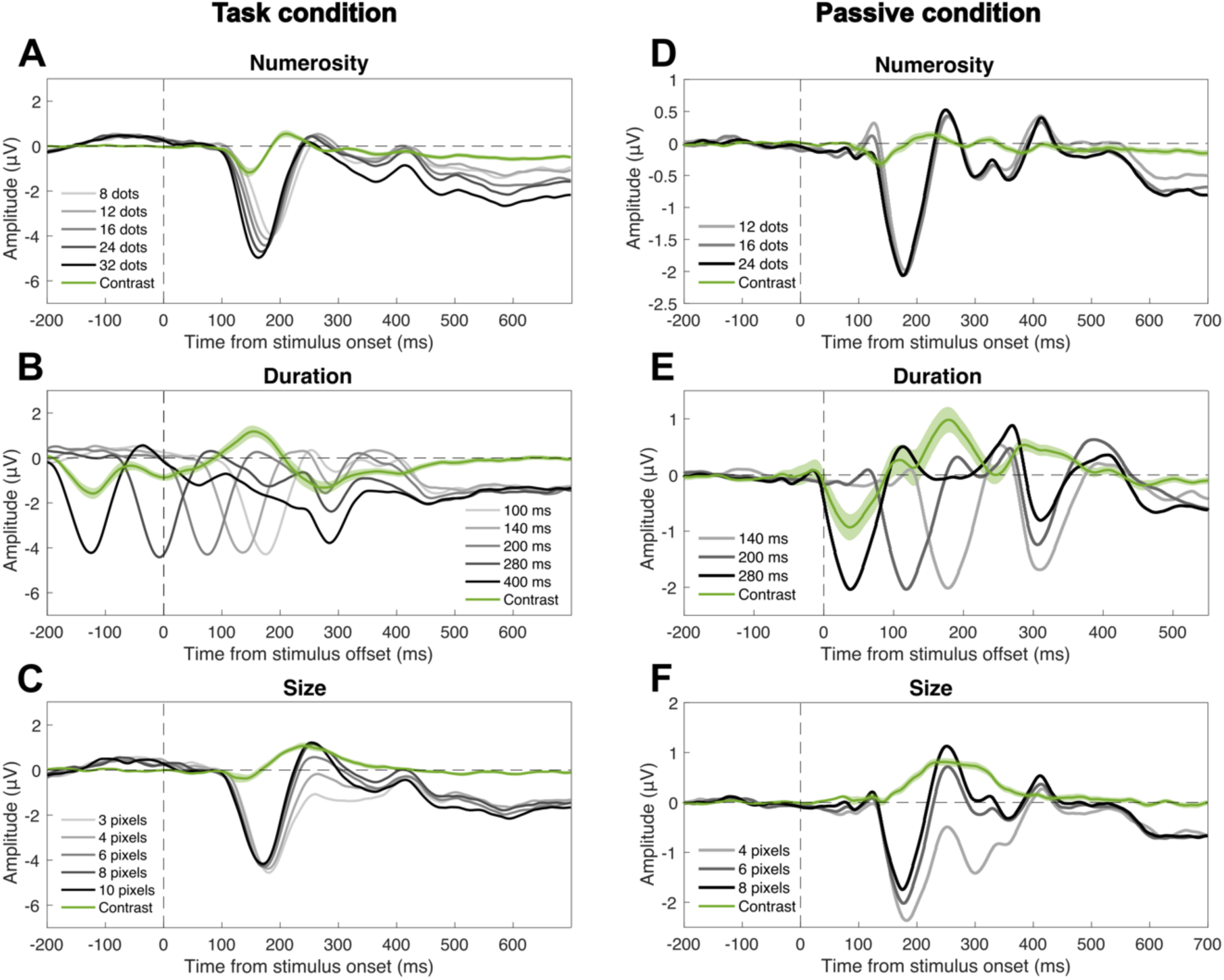
Event-related potentials (ERPs) evoked by each magnitude in the two experimental conditions. Panels A-C: data from the task condition. Panels D-F: data from the passive condition. (A) ERPs evoked by the stimulus numerosity in the task condition. (B) ERPs evoked by the stimulus duration in the task condition. (C) ERPs evoked by item size in the task condition. (D) ERPs evoked by the stimulus numerosity in the passive condition. (E) ERPs evoked by the stimulus duration in the passive condition. (F) ERPs evoked by item size in the passive condition. In all panels, the green wave indicates the linear contrast of the ERPs. Note that while numerosity and size ERPs were time-locked to the onset of the stimuli, ERPs corresponding to duration were re-aligned to the offset. The zero in panel B and E thus indicates the offset of the stimuli. The vertical dashed line indicates the onset or offset of the stimuli. The horizontal dashed line indicates the zero of the amplitude scale. All the ERPs are the average of signals from channels Oz, O1, and O2. The shaded area around the green contrast wave represents the SEM.

To better address the significance of magnitude-sensitive brain responses, we computed a measure of ERP contrast based on the difference between the extreme levels of the interfering dimensions’ ranges. Namely, for each level of each magnitude, we contrasted the ERPs as a function of the extreme levels of the interfering dimensions. For example, to compute the effect of numerosity, we subtracted the ERP corresponding to the combination of 100 ms and the two extreme levels of numerosity (8 and 32 dots). The same subtraction was performed for the combination of 140, 200, 280, and 400 ms and the extreme levels of numerosity. The same was done for the combination of each level of size and the extreme levels of numerosity. The effect of duration and size were computed in the same way by switching the dimension. The average of this contrast measure, reflecting the effect of the different magnitudes in driving ERPs, is shown in Fig. 4. Additionally, Fig. 4 shows the topographic distribution of contrast amplitude in a 50-ms window around the main peak of each corresponding contrast wave. To assess the significance of the contrast amplitude, we performed a series of one-sample t-tests against zero, corrected for multiple comparisons with false discovery rate (FDR; q = 0.05). When reporting the results below, we indicate the range of t-values and FDR-adjusted p-values as [min max].

**FIGURE 4.**
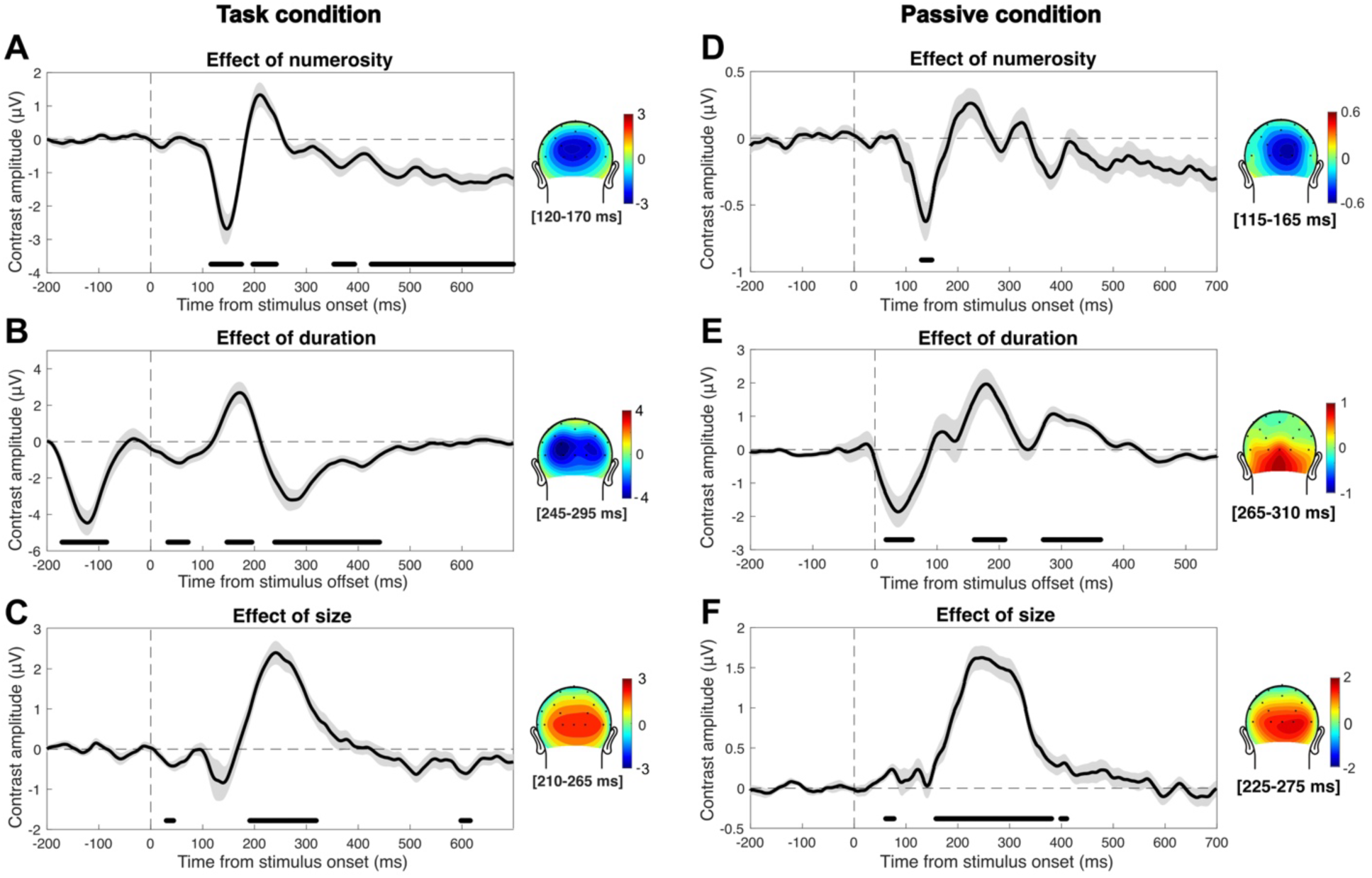
Average contrast amplitude reflecting the neural effects of the three magnitudes. Panels A-C: data from the task condition. Panels D-F: data from the passive condition. The contrast amplitude in this case was computed as the difference between the extreme levels of each “interfering” magnitude, separately for each level of each magnitude. For instance, the effect of numerosity on duration was computed as the difference between ERPs corresponding to 100 ms/32 dots and 100 ms/8 dots, and so on for the other levels of duration. The same was done for the effect of numerosity on size, and the resulting contrasts were averaged together to increase the signal-to-noise ratio. A similar procedure was then used to compute the effect of duration and the effect of size. (A) Contrast amplitude reflecting the effect of numerosity in the task condition. (B) Contrast amplitude reflecting the effect of duration in the task condition. (C) Contrast amplitude reflecting the effect of size in the task condition. (D) Contrast amplitude reflecting the effect of numerosity in the passive condition. (E) Contrast amplitude reflecting the effect of duration in the passive condition. (F) Contrast amplitude reflecting the effect of size in the passive condition. The black lines at the bottom of the plots mark the significant latency windows assessed with a series of FDR-corrected one-sample t-tests. The vertical dashed line indicates the onset or offset of the stimuli. The horizontal dashed line indicates the zero of the amplitude scale. The shaded area around the wave represents the SEM. The topographic plots besides each panel show the distribution of scalp activity in a 50-ms window around the main peak of each wave. All waves shown in the figure reflect the average of signals from channels Oz, O1, and O2.

The numerosity-sensitive brain responses (Fig. 4A) showed four significant latency windows. The strongest effect was observed at a negative deflection at 120-175 ms (t = [-6.97, -2.41], p = [<0.001, 0.049]) after stimulus onset, which was around the peak of contrast amplitude (-2.7 μV) observed at 145 ms after stimulus onset. This peak was followed by additional significant windows at 200-240 ms (t = [2.45, 3.62], p = [<0.001, 0.046]), 355-390 ms (t = [-3.18, -2.41], p = [0.012, 0.048]), and 425-700 ms (t = [-5.61, -2.47], p = [0.001, 0.044]) after stimulus onset. Regarding the effect of duration on ERPs, we observed four significant latency windows. The first one was observed before stimulus offset, spanning from -170 to -85 ms (t = [-6.55, -2.58], p = [<0.001, 0.049]). Then we observed two relatively early windows at 30-70 ms (t = [-3.66, -2.58], p = [0.007, 0.049]) and 145-195 ms (t = [2.58, 4.50], p = [0.003, 0.049]). Looking at the ERPs shown in Fig. 3B, these three latency windows however appear to be driven each by a single duration level, due to the onset responses (i.e., only one wave shows a deflection while the others are flat). Such responses cannot thus be considered as genuine correlates of duration, but are spurious effects due to the re-alignment of brain waves to the offset of the stimuli. A more genuine peak of activity driven by duration was instead observed at 270 ms after the offset (-3.2 μV), and we observed a significant latency window around this peak, spanning 240-440 ms (t = [-5.99, -2.60], p = [<0.001, 0.048]). In the case of size, the peak of activity was observed at 240 ms (2.4 μV). The largest significant latency window was observed around this peak, spanning 190-320 ms (t = [3.14, 8.44], p = [<0.001, 0.049]). Two additional, smaller significant windows were observed at very early latencies (30-45 ms; t = [-4.04, -3.23], p = [0.009, 0.042]), and at later latencies (600-615 ms; t = [-3.59, -3.16], p = [0.022, 0.048]). In all cases, the topographic distribution of scalp activity around the peaks (plotted besides each panel, Fig. 4A-C) showed a posterior distribution consistent with activity in the occipital cortex.

Our main goal in the task condition was however to identify the latency windows whereby the modulation of brain activity predicts the magnitude integration bias observed behaviorally. We then further computed two measures of the effect that could be related to each other in data analysis: ΔPSE, reflecting the behavioral effect, and ΔERP, reflecting the neural effect of magnitudes. These measures were computed by subtracting either the PSE or the ERP amplitude of each level of the interfering magnitudes from the PSE/ERP corresponding to the middle, reference level. For the ΔERP, this measure was computed at each time point throughout the epochs, and separately for the different types of task (see Fig. 5). Doing so, we thus computed the effect of numerosity in the duration and size task (Fig. 5A-B), the effect of duration in the numerosity and size task (Fig. 5C-D), and the effect of size in the numerosity and duration task (Fig. 5E-F).

**FIGURE 5.**
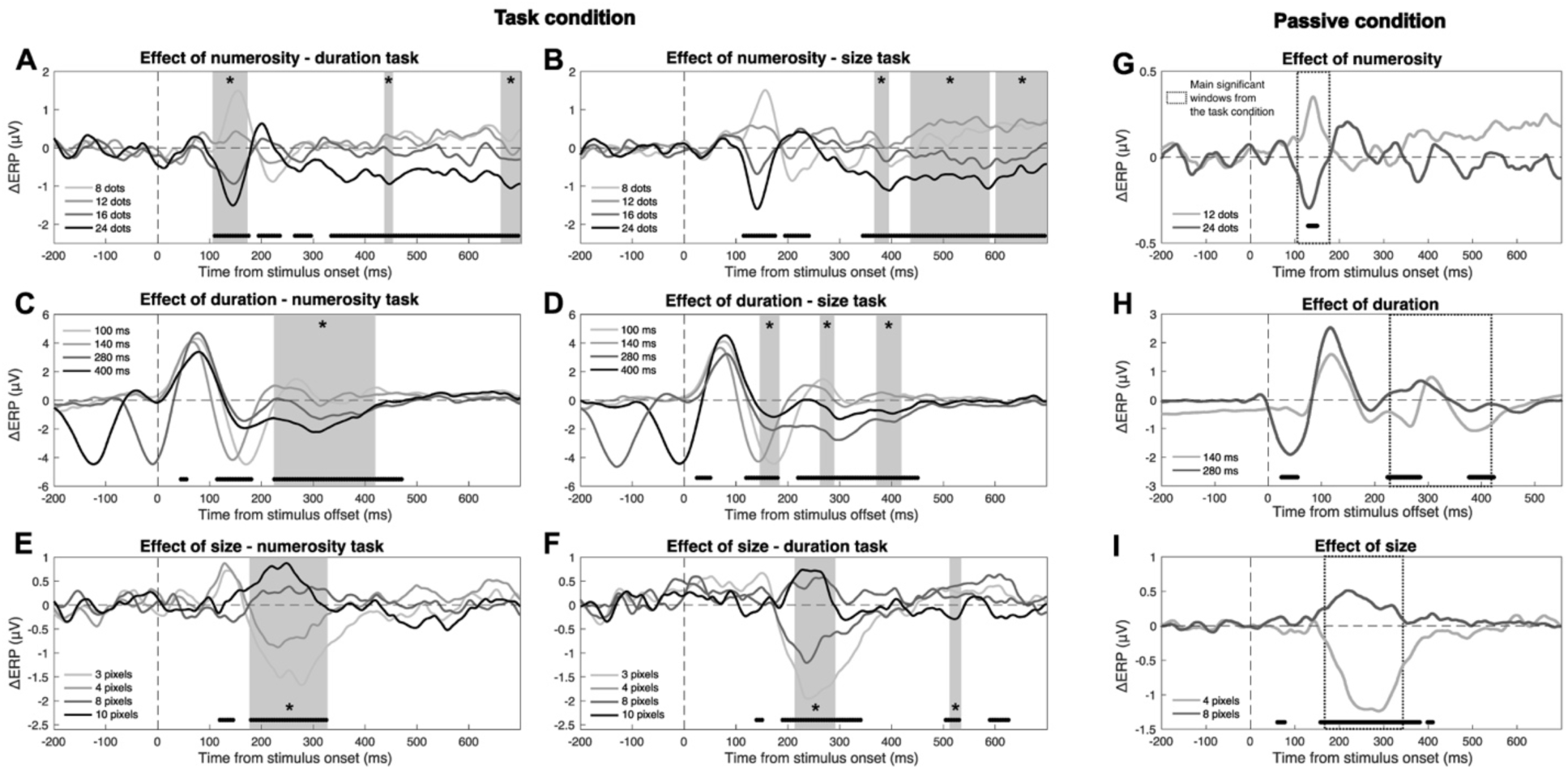
ΔERP measures and relationship with the behavioral effect. Panels A-F: data from the task condition. Panels G-I: data from the passive condition. (A) ΔERP measures reflecting the effect of numerosity in the duration task. (B) Effect of numerosity in the size task. (C) Effect of duration in the numerosity task. (D) Effect of duration in the size task. (E) Effect of size in the numerosity task. (F) Effect of size in the duration task. (G) ΔERP measures reflecting the effect of numerosity in the passive condition. (H) Effect of duration in the passive condition. (I) Effect of size in the passive condition. The black lines at the bottom of the plots mark the latency window where we observed a significant difference in ΔERP as a function of the different levels of the magnitude. The vertical dashed line indicates the onset or offset of the stimuli. The horizontal dashed line indicates the zero of the amplitude scale. The grey shaded areas marked with stars indicate the latency windows where we observed a significant relationship between ΔERP and the behavioral effect (ΔPSE). The dotted boxes in panels G-I (passive condition) show the main latency windows whereby the brain responses in the task condition predicted the behavioral effect. All waves shown in the figure reflect the average of signals from channels Oz, O1, and O2.

To address the relationship between neural and behavioral measures of magnitude effects, we first looked for latency windows showing a significant modulation of ΔERP as a function of the different levels of the magnitudes. To do so, we performed a series of LME tests individually for the effect of each magnitude in each task. In the LME model, we entered the ΔERP as the dependent variable, the ratio of each magnitude level with the middle level as the predictor, and the subjects as the random effect. The LME tests were performed across a series of 10-ms windows with a 5-ms step, in a sliding-window average fashion. To control for multiple comparisons, we again applied a FDR procedure with q = 0.05. Clusters of less than three consecutive significant tests (after FDR) were not considered. The results of these tests are shown with black lines at the bottom of each plot in Fig. 5, marking the significant latency windows.

After identifying the latencies showing a significant modulation of ΔERP, we looked for a relationship between ΔERP and ΔPSE within these windows. We thus performed a series of LME tests (10-ms windows with 5-ms step) including ΔPSE as the dependent variable, ΔERP as the predictor, and the subjects as the random effect. The effect of numerosity on duration (Fig. 5A) showed three windows whereby the modulation of ΔERP could predict the behavioral effect (marked with grey shaded areas in the figure), a larger early window at 110-170 ms, followed by two smaller windows at 440-450 ms and 665-695 ms (β = [0.006, 0.009], t = [2.02, 3.69], p = [<0.001, 0.047], adj-R^2^ = [0.31, 0.43]). The effect of numerosity on size (Fig. 5B) showed again three significant windows, but clustered at later latencies: 370-390 ms, 440-585 ms, and 605-695 ms (β = [0.085, 0.180], t = [2.06, 4.12], p = [<0.001, 0.047], adj-R^2^ = [0.25, 0.36]). The effect of duration on numerosity showed a single large significant window, spanning 225-415 ms (β = [0.233, 0.597], t = [2.20, 3.77], p = [<0.001, 0.31], adj-R^2^ = [0.47, 0.54]). The effect of duration on size, showed three smaller windows, with the first at earlier latencies spanning 150-180 ms, and the following two at latencies more consistent with the effect on numerosity: 265-285 ms and 375-415 ms (β = [-0.043, 0.088], t = [-2.67, 2.78], p = [0.006, 0.047], adj-R^2^ = [0.56, 0.59]). Finally, the effect of size on numerosity showed a single, large significant window at 180-325 ms (β = [0.707, 1.171], t = [2.65, 6.50], p = [<0.001, 0.009], adj-R^2^ = [0.18, 0.48]), while the effect of size on duration showed two significant windows at 215-290 ms and 515-530 ms (β = [-0.012, 0.013], t = [-3.19, 2.12], p = [0.002, 0.041], adj-R^2^ = [0.13, 0.26]). These results show that the behavioral effect of magnitude integration could be reliably predicted by the modulation of magnitude-sensitive responses in the different task types, providing a neural signature of the effect.

### Passive condition

In the passive-viewing condition, participants watched a stream of dot-array stimuli modulated in numerosity, duration, and item size, and responded to occasional oddball stimuli defined by a lower contrast. No instruction suggested the participants to explicitly attend the magnitudes of the stimuli. This passive-viewing protocol thus provides a cleaner index of the responses to the different magnitudes, not confounded by decision making or other task-related processes. If magnitude integration arises from automatic perceptual processes, then we expected to observe a similar modulation of brain responses consistent with the timing observed in the task condition (see Fig. 5). Instead, if magnitude-related decision making is necessary for magnitude integration to occur, the modulation of brain responses linked to the behavioral effect should not occur during passive viewing.

First, we assessed the ERPs corresponding to the different levels of the three magnitudes (Fig. 3D-F). The overall pattern was largely consistent with what observed in the task condition (see Fig. 3A-C), with however some differences. Numerosity (Fig. 3D) showed an early positive deflection that we did not observe in the task condition, with the magnitude of the stimuli however modulating the stimuli negatively (i.e., the smaller the numerosity, the higher the positive deflection in response amplitude). Additionally, ERPs at later latencies showed a weaker modulation compared to the task condition. Duration (Fig. 3E) showed a similar deflection compared to the task condition, but the modulation of amplitude was in the opposite direction (i.e., see topographic plots besides the panels). Finally, size (Fig. 3F) showed instead ERPs consistent with the task condition.

To better assess the significance of the magnitude-sensitive brain responses, we computed again a measure of contrast based on the difference between the extreme levels of each magnitude (Fig. 4D-F), as in the task condition. The contrast amplitude was then tested with a series of one-sample t-tests against zero, corrected with FDR (q = 0.05). In the case of numerosity (Fig. 4D), we observed a significant early window (130-150 ms; t = [-4.34, -3.64], p = [0.026, 0.047]), showing a negative deflection consistent with the task condition (see Fig. 4A). The peak of activity in this window (-0.62 μV) was at 140 ms. Differently from the task condition, we did not observe significant latency windows later on in the epoch. The effect of duration (Fig. 4E) showed three significant windows. The first one at 20-60 ms (t = [-3.99, -2.68], p = [0.003, 0.049]), the second at 160-210 ms (t = [2.68, 4.34], p = [0.002, 0.049]), and the third at 270-360 ms (t = [2.71, 4.76], p = [0.002, 0.043]). Note however that similarly to the task condition, the first two significant windows appear to be mostly driven by the onset responses of individual durations, while the last window shows a consistent deflection in responses corresponding to all the different levels of duration (see Fig. 3E). The topographic plot showing the distribution of peak activity (Fig. 4E) thus reflects this last latency window (peak at 290 ms, 1.05 μV). Differently from the task condition, however, the contrast amplitude here showed a positive, rather than negative, deflection. Finally, size (Fig. 4F) showed a large main window at 155-380 ms (t = [2.62, 10.88], p = [<0.001, 0.048]), with a peak at 245 ms (1.62 μV) consistent with the effect of size in the task condition. In addition, we observed two additional, smaller windows at 60-75 ms and 400-410 ms (t = [2.62, 3.63], p = [0.005, 0.049]). Similarly to the task condition, the topography of peak amplitude over the scalp showed a posterior distribution consistent with occipital cortex (see plots beside panel D-F).

In order to better compare the modulation of brain responses in the task and in the passive condition, we also computed the ΔERP measure (Fig. 5G-I). The ΔERPs were assessed with a series of paired t-tests, performed considering 10-ms windows (step = 5 ms) as in the task condition, corrected with FDR (q = 0.05). The overall timing of the significant windows was consistent with the contrast measure (see Fig. 4D-F). ΔERPs reflecting the effect of numerosity (Fig. 5G) showed a significant modulation at 130-150 ms (t = [3.64, 4.33], p = [0.025, 0.047]). The effect of duration (Fig. 5H) showed a modulation at 225-285 ms (t = [-6.91, -2.78], p = [<0.001, 0.049]), and at 380-420 ms (t = [-3.68, -2.79], p = [0.016, 0.049]) after stimulus offset. Finally, in the case of size (Fig. 5I), we observed the main modulation in a large window spanning 160-380 ms (t = [-10.87, -2.62], p = [<0.001, 0.049]). We again observed two smaller significant windows at 60-75 ms (t = [-3.63, -2.65], p = [0.005, 0.046]) and 400-410 ms (t = [-2.85, -2.62], p = [0.032, 0.049]). As a comparison, Fig. 5G-I shows with dotted boxes the latency windows where we observed a significant relationship between neural and behavioral measures of magnitude integration in the task condition (Fig. 5A-F). In all cases, we observed an overlap between the significant windows in the passive and task condition.

### Multivariate decoding analysis

To achieve a more direct comparison of magnitude-sensitive brain activity during the task and in passive viewing, we performed a multivariate “decoding” analysis across the two experimental conditions. In the analysis, we trained a classifier (support vector machine) with data from the task condition, and tested its ability to decode magnitude-sensitive brain activity in the passive condition. This training and testing direction was chosen to obtain a larger set of classification accuracy (CA) values, in order to achieve more robust and stable results when testing the statistical significance of the decoding. Indeed, the analysis was performed by training the classifier on a set of data points each formed by the average data of one participant. The classifier was then tested separately on the average data of each participant in the passive condition group, according to a leave-two-out procedure, i.e., two datapoints each corresponding to a class entered into the analysis were left out from the training set, and two independent datapoints from the passive conditions were used for testing. This procedure thus resulted in a distribution of CA values corresponding to the number of participants in the passive condition. We did not run the analysis in the opposite direction (i.e., training with the passive data and testing with the task data) since, due to the different number of data points (i.e., due to the different number of participants in the two conditions), the results would be difficult to combine. See *Methods* for more information about the decoding procedure. According to our hypothesis, if the brain responses related to magnitude processing and integration are similar irrespective of the task, then the classifier should be able to decode magnitude information from the passive data. Otherwise, if magnitude processing entails mechanisms specific to the task performed, no above-chance decoding should be observable. The ability of the classifier to decode magnitude information was evaluated based on the distribution of CA values, obtained across a series of small time windows (i.e., 15-ms window with 5-ms step) throughout the epochs. The distribution of CA values at each time window was then tested with a permutation (sign flipping) test to assess whether it resulted significantly higher than chance level (0.5; see Methods for more information). The results of this analysis are shown in Fig. 6.

**FIGURE 6.**
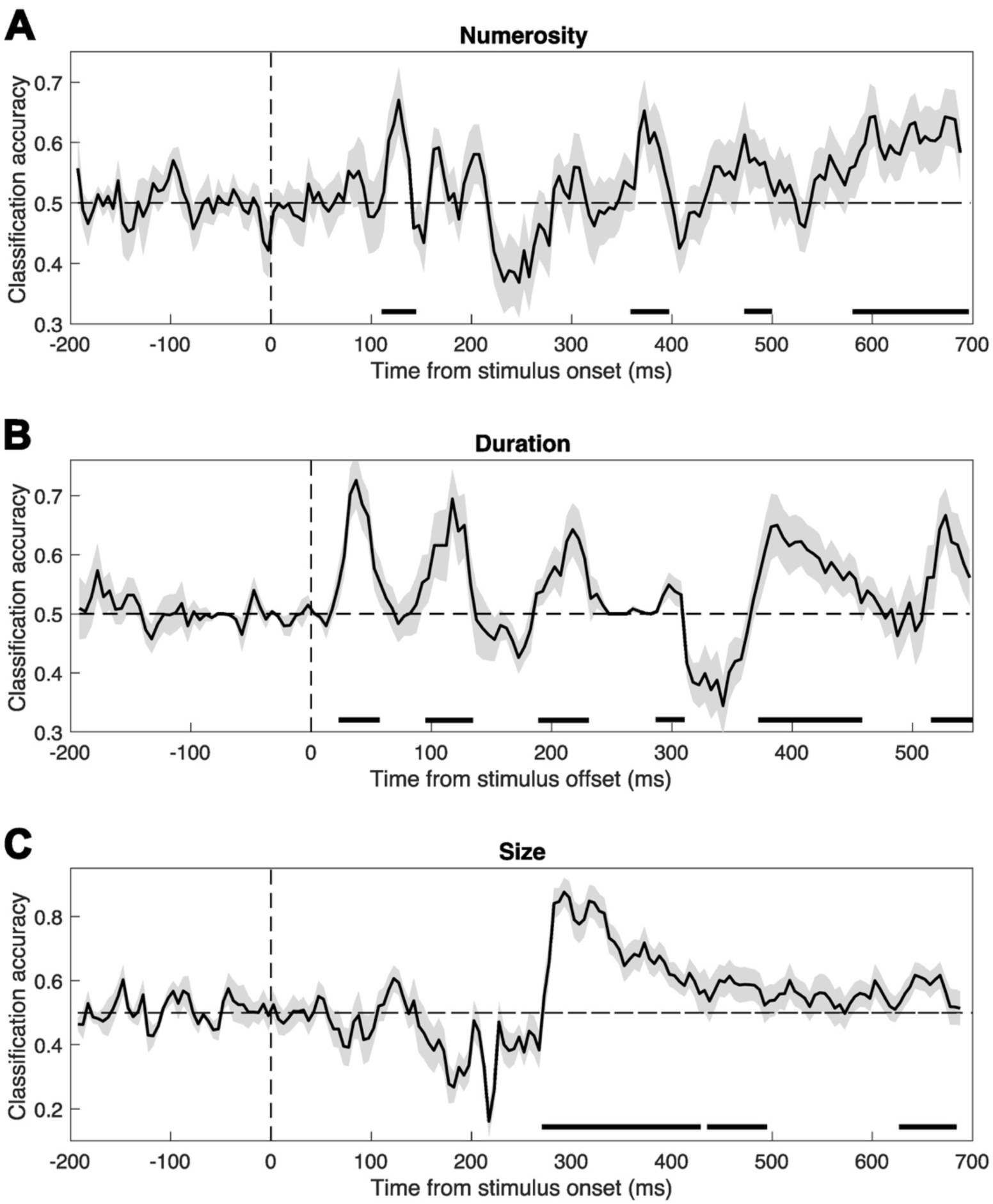
Results of the multivariate decoding analysis. The decoding analysis was performed by training a classifier on data from the task condition, then tested on data from the passive condition, in order to achieve a more direct comparison of the magnitude-related brain processes in the two experimental conditions. (A) Classification accuracies obtained in the decoding of numerosity. (B) Classification accuracies obtained in the decoding of duration. (C) Classification accuracies obtained in the decoding of size. The horizontal dashed lines indicate the chance level (0.5). The vertical dashed lines mark either the time of stimulus onset for numerosity and size, or the time of stimulus offset, for duration. The shaded area around the waves indicates the SEM, which represents the variability across the distribution of classification accuracy values obtained in the decoding procedure. The black lines at the bottom of the plot mark the latency windows where the decoding is significantly above-chance, as observed with a series of permutation tests.

Overall, the multivariate analysis revealed several latency windows in which magnitude-sensitive brain responses in the passive condition could be successfully predicted based on the training with the task condition data. Below, the results are reported in terms of the range of CAs observed (CA = [min, max]) and p-values of the permutation tests (p = [min, max]). In the case of numerosity, the analysis showed significant above-chance decoding at four latency windows. Namely, an early window spanning 110-145 ms after stimulus onset (CA = [0.57, 0.67], p = [<0.001, 0.010]), followed by later windows at 360-395 ms (CA = [0.59, 0.65], p = [<0.001, 0.049]), 475-500 ms (CA = [0.56, 0.57], p = [0.026, 0.049]), and 585-695 ms (CA = [0.57, 0.64], p = [<0.001, 0.036]). In the case of duration, we observed six significant latency windows, at 20-60 ms after stimulus offset (CA = [0.58, 0.73], p = [<0.001, 0.013]), 95-135 ms (CA = [0.62, 0.69], p = [<0.001, 0.017]), 185-235 ms (CA = [0.54, 0.64], p = [<0.001, 0.038]), 285-310 ms (CA = [0.53, 0.55], p = [0.004, 0.048]), 370-460 ms (CA = [0.56, 0.65], p = [<0.001, 0.039]), and 515-550 ms (CA = [0.58, 0.67], p = [<0.001, 0.035]). Finally, in the case of size, we observed three significant latency windows, at 270-430 ms (CA = [0.60, 0.85], p = [<0.001, 0.020]), 440-495 ms (CA = [0.58, 0.61], p = [0.008, 0.047]), and 630-680 ms (CA = [0.58, 0.62], p = [<0.001, 0.049]).

## DISCUSSION

In the present study, we assessed and compared the signatures of magnitude integration in two different conditions: when participants are engaged in actively judging the magnitude of the stimuli, or when they passively watched the stimuli. The phenomenon of magnitude integration – i.e., the mutual biases usually observed across different dimensions – is a hallmark of magnitude perception. Indeed, stimulus dimensions such as numerosity, duration, and size systematically interact with each other, leading to biases when judging them. Such mutual interactions have played a pivotal role in the development of influential theories like “a theory of magnitude” (ATOM; Walsh, 2003) and the “metaphor” theory (Casasanto & Boroditsky, 2008). However, the nature of this bias and its underlying mechanisms remain unclear.

Different mechanisms have been proposed to explain the interaction of magnitude dimensions. On the one hand, according to ATOM, the interaction would occur in perceptual processing due to the encoding of different dimensions with a common neural code (Walsh, 2003). In support of such a perceptual account of magnitude integration, we have recently shown that the effect relies on a mechanism similar to perceptual binding, inducing a positive bias across dimensions only when they are conveyed by the same stimulus (i.e., as opposed to magnitudes conveyed by separate, superimposed stimuli; Togoli, Bueti, et al., 2022). Recent neuroimaging studies, however, albeit showing common neural substrates, failed to provide evidence for a shared neural code (Borghesani et al., 2019; Hendrikx et al., 2024; Tsouli et al., 2022). According to the metaphor theory, on the other hand, the effect would instead arise at the conceptual or linguistic level, due to the use of “spatial” concepts to describe time (e.g., a “long” time; Bottini & Casasanto, 2013; Casasanto & Boroditsky, 2008; but see Whitaker et al., 2022). This theory however relies on asymmetric effects across temporal and non-temporal dimensions, which depend on the type of stimuli used (Javadi & Aichelburg, 2012; Lambrechts et al., 2013; Togoli et al., 2021). Moreover, other authors proposed that magnitudes could interact during working memory maintenance, nudging each other while stored in memory (Cai et al., 2018; Cui et al., 2022), or bias the response selection in comparisons tasks (Yates et al., 2012). Considering the results from these studies, whether magnitude integration across dimensions (e.g., numerosity, duration, and size) occurs at a perceptual or at a post-perceptual stage remains a debated topic.

In the present study, we further addressed the nature of the magnitude integration effect by assessing a new prediction. Namely, a high-level effect hinging upon magnitudes concurrently held in memory (i.e., one magnitude biasing the memory of the other) or on active decision-making (i.e., one magnitude interfering with the response to another magnitude) should show a unique neural signature not present when the magnitudes are neither explicitly attended nor judged. Conversely, a perceptual effect is expected to occur in a more automatic fashion, independently from the relevance or judgment of magnitude. Thus, similar signatures should be observable with or without a magnitude judgment task.

Our behavioral results show systematic biases across the three magnitudes. First, numerosity was biased by both duration and item size. However, while duration show a congruent effect (the longer the duration, the higher the perceived numerosity) as in previous studies (Javadi & Aichelburg, 2012; Togoli et al., 2021), size induces an opposite bias. Although different from the relationship between other dimensions, previous studies indeed show that the effect of dot size on numerosity entails a negative effect, so that the larger the dot size, the lower the perceived numerosity (DeWind et al., 2015; Fornaciai et al., 2019). Duration is instead similarly affected by both numerosity and size in a congruent fashion, in line with previous studies (e.g., Lambrechts et al., 2013; Xuan et al., 2007), although the latter exerts a stronger influence. Finally, size seems the dimension most resistant to integration biases, and shows only modest, albeit significant, influences from the other magnitudes. Size is also the dimension that is easiest to judge (Fig. 2D), and the generally lower variability of responses might explain its robustness to biases. However, in a previous study from our group addressing trial-history effects in different magnitude dimensions (i.e., “serial dependence” effects; Fornaciai et al., 2023), size showed stronger biases compared to duration and numerosity, while again showing the highest precision. Thus, the perception of size does not seem intrinsically more resistant to biases, and the lower effect observed here might be a feature of magnitude integration effects rather than a general property. Considering the pattern of effects across dimensions, the results thus show some partial asymmetries, as some dimensions are more vulnerable to biases than others, in line with previous studies using similar stimuli (Togoli, Bueti, et al., 2022; Togoli et al., 2021).

In terms of event-related potentials, in the task condition we found robust brain responses to the different magnitudes. Overall, our analyses identified a set of latency windows that show the stronger peaks of activity driven by the different dimensions. Namely, around 150 ms and 250 ms after stimulus onset in the case of numerosity and size, and around 300 ms after stimulus offset for duration. Brain activity at these latency windows appears to be modulated by the different dimensions in a parametric fashion, according to the magnitude of the stimuli. Crucially, with just one exception (i.e., the effect of numerosity in the size task), brain activity at or around such peaks can significantly predict the bias observed behaviorally: the larger the brain responses, the stronger the magnitude integration bias. In the case of numerosity, the timing observed here (∼150 ms) is consistent with numerosity-sensitive responses measured in previous studies. Although this timing is slightly earlier compared to the P2p component (∼200 ms), i.e., the ERP component most often associated with numerosity (Grasso et al., 2022; Libertus et al., 2007; Park et al., 2016; Temple & Posner, 1998), several studies also showed numerosity-sensitive responses at earlier latencies, starting at around 75-100 ms after stimulus onset (Fornaciai et al., 2017; Fornaciai & Park, 2018; Park et al., 2016). In the case of duration, previous studies highlight a variety of possible EEG correlates of duration processing, like the contingent negative variation (CNV; Damsma et al., 2021; but see Kononowicz & Penney, 2016), the N2 (Tonoyan et al., 2022), the P2 (Li et al., 2017), and the P3 (Ernst et al., 2017) ERP components. The timing shown in our results appears to be consistent with the results of Benau et al. (2018), showing duration sensitivity at around 350 ms after stimulus offset. Finally, in terms of size, the timing of responses sensitive to the size of the items appears to be roughly consistent with previous results (Park et al., 2016) showing a peak at around 200 ms.

The timing of magnitude-sensitive brain response in passive viewing revealed similar evoked activity in most of the cases, closely mirroring the responses observed in the task condition. Especially in the case of numerosity and size, the peaks of magnitude-sensitive activity (i.e., ΔERP; compare Fig. 5A-F with Fig. 5G-I) show indeed a one-to-one correspondence, with similar timing and polarity. In duration perception, however, although the timing and topography of responses is very similar, we observed ERPs with an opposite polarity. This may additionally suggest that while the processing of numerosity and size is largely invariant across the two conditions, the brain responses to duration may at least partially depend on the task relevance of this dimension. Namely, while the same duration processing stage seems to get engaged (i.e., as suggested by brain responses at the same latency and with the same scalp topography), actively attending the magnitudes of the stimuli may modulate how duration information is processed. This is not completely surprising, as duration shows different properties compared to the other dimensions (i.e., duration information needs to be accumulated, while the other dimensions can be processed from the onset), and the encoding of duration information is notoriously poorer in vision compared to other senses (e.g., Alais & Burr, 2004; Cai & Connell, 2015). Thus, the processing of duration may be more sensitive to the modulatory effects of attention or task-relevance.

The lack of decision-making processes related to magnitude in the passive viewing parading represents its major strength, as it allows to exclude the involvement of task-related brain processes (e.g., working memory encoding and maintenance of magnitude information; Cai et al., 2018; Cui et al., 2022), and response biases (Yates et al., 2012). However, it also has the obvious weakness that magnitude integration could not be directly measured to confirm the effect. The striking similarity in the brain responses to the magnitudes, peaking at the same latencies where we demonstrated a relationship with the behavioral effect and showing the same scalp topography, nevertheless provides evidence that magnitude integration likely occurs even in the absence of a task. While this comparison remains qualitative, the multivariate “cross-condition” decoding analysis provides quantitative evidence that the brain activity at several latency windows does not depend on the presence of a magnitude task. Indeed, the ability of the classifier to successfully decode the brain responses to magnitude across conditions shows that similar brain processes are engaged at specific time points, resulting in similar patterns of brain activity. In all cases, the latency windows showing above-chance decoding are largely consistent with the most important windows highlighted in the other analyses (e.g., in terms of ΔERP and its relationship with the behavioral effect). Namely, the timing of above-chance decoding in the case of numerosity (i.e., the 110-145 ms window), duration (i.e., 370-460 ms), and size (i.e., 270-430 ms) overlaps with similar windows showing a relationship between ERPs and behavior (in the task condition analysis), and with the effect of the different magnitudes in the passive condition. The analysis also highlights several other windows showing significant cross-condition decoding, suggesting that similar patterns of brain activity emerge at multiple processing stages across conditions, both early and late. The decoding analysis thus provides further evidence that magnitude processing entails patterns of brain activity largely independent from the task. In other words, with this analysis we demonstrate that the task and passive condition not only entail similar magnitude-sensitive responses at the same processing stages, but also that such responses show very similar patterns of activity likely reflecting the same brain processes. In turn, this also suggest that the magnitude integration phenomenon – which is reflected by activity at such latency windows – likely takes place irrespective from the task, in an automatic and perceptually-driven fashion. According to this interpretation, the integration of different magnitudes and the relative bias would occur via perceptual processes affecting how we experience the different dimensions, and not only their memory traces or how they are judged. Namely, for instance, when we underestimate a duration because it is paired with a low numerosity we perceptually experience a shorter duration.

Differently from the present study, previous EEG results concerning magnitude integration (involving duration and length) suggested the involvement of working memory interference (Cui et al., 2022). Cui et al. (2022) indeed observed effects of duration and length after the offset of the intervals, at ERP components usually associated with working memory maintenance, such as the P2 and P3b. While the effect of duration in Cui et al.’s work shows a timing consistent with the present results (∼250-300 ms after stimulus offset), length has an effect at a much different timing (∼300 ms after stimulus *offset*) compared to our earlier peak of responses to the size dimension (∼250 ms after stimulus *onset*). Additionally, the scalp topography of the magnitude effects had a much more anterior distribution, peaking at parieto-frontal locations, as opposed to our results showing a posterior, occipital distribution. However, Cui et al. also employed much different stimuli: longer intervals (800-1,200 ms) and quite large lengths up to 15 degrees of visual angle. Considering the relatively long durations and the fact that the stimuli were marked only at the beginning (onset) and end (offset) point, it is not surprising that they engaged memory processes (e.g., see for instance Rammsayer & Lima, 1991). Both magnitudes in such a task indeed rely on the memory trace of the first marker presented rather than on sustained sensory stimulation, making it more likely that any interference would involve higher-level memory processes. Our results however do not conflict with such an interpretation. Perceptual and mnemonic interferences are indeed not mutually exclusive processes, and can both occur depending on the nature of the stimuli and paradigm used. Our results however show that magnitude interactions can be perceptual in nature when based on stimuli relying on sensory/perceptual processing rather than memory.

How would this perceptual interaction occur in the brain? Interestingly, our results show that the effects across the different magnitudes do not occur at a unique, generalized stage, but show different timings consistent with different brain processing stages. The interference between magnitudes thus depends on the specific processing dynamics of the different dimensions (see also Togoli et al., 2021), rather than a common processing stage. Our results are thus not fully consistent with the idea of a generalized magnitude processing system, as proposed by the ATOM framework (e.g., Walsh, 2003). Instead, the results seem more in line with recent findings of separate topographical cortical maps of different magnitudes, partially overlapping with each other (Fortunato et al., 2023; Harvey et al., 2013, 2015; Harvey & Dumoulin, 2017; Hendrikx et al., 2024; Protopapa et al., 2019). Recently, it has indeed been proposed that the interaction between different magnitudes could arise from the overlap of neural populations sensitive to different dimensions but without neural alignments across dimensions (Hendrikx et al., 2024; Tsouli et al., 2022), therefore arguing against the existence of a centralized mechanism or a common magnitude neural code (Walsh, 2003).

To conclude, our results show that the neural signatures of magnitude processing and integration are very similar whether participants explicitly attend and judge magnitude information or passively watch the stimuli. This in turn suggests that similar brain processing stages are engaged irrespective of the task, and thus that magnitude integration likely occur even in the absence of magnitude decision-making. Overall, our results thus provide new evidence supporting the idea that magnitude integration is an automatic perceptual phenomenon independent from the task performed, affecting the phenomenological appearance of the stimuli rather than their memory or the way they are judged.

## Data and code availability

All the data generated in the experiments described in this manuscript and the experimental code are available on Open Science Framework, at this link: https://osf.io/sn9h8/

## Acknowledgements

This project has received funding from the European Union’s Horizon Europe research and innovation programme under the Marie Sklodowska-Curie grant agreement No. 101103020 “PreVis” to MF, from the European Research Council (ERC) under the European Union’s Horizon 2020 research and innovation programme grant agreement No. 682117 BIT-ERC-2015-CoG to DB, and from the Italian Ministry of University and Research under the call FARE (project ID: R16X32NALR) and under the call PRIN2017 (project ID: XBJN4F) to DB. IT is supported by a FSR incoming post-doctoral fellowship granted by Université Catholique de Louvain. OC is a senior research associate at the National Fund for Scientific Research (FRS-FNRS)-Belgium.

## Author contributions

IT: conceptualization, methodology, software, investigation, formal analysis, data curation, visualization, writing – original manuscript, writing – review and editing. OC: conceptualization, resources, writing – original manuscript, writing – review and editing. DB: resources, funding acquisition, writing – review and editing. MF: conceptualization, methodology, formal analysis, visualization, supervision, project administration, funding acquisition, writing – original manuscript, writing – review and editing.

## Competing interests

The authors declare no competing interest.

## Notes

https://osf.io/sn9h8/

### Competing Interest Statement

The authors have declared no competing interest.

### Summary of Updates

- Added multivariate decoding analysis to better compare the magnitude-sensitive brain responses across the two experimental conditions. - Revised text to improve the writing

